# A rodent model of social rejection

**DOI:** 10.1101/066993

**Authors:** Haozhe Shan, Inbal Ben-Ami Bartal, Peggy Mason

**Affiliations:** Department of Neurobiology, University of Chicago, Chicago, United States

**Author notes:** Current address: 188 Li Ka Shing Center, Berkeley CA 94720. The authors declare no competing interests. Contributions: HZS, Conception and design, Acquisition of data, Analysis and interpretation of data, Drafting or revising the article, IB-AB, Conception and design, Drafting or revising the article, PM, Conception and design, Analysis and interpretation of data, Drafting or revising the article.

**Keywords:** Apathy, Prosocial behavior, social rejection, helping, anxiety

## Abstract

Rats help conspecifics by releasing them from a restrainer only when they have previous social experience with the strain of the restrained rat. When rats have been cross-fostered with rats of a different strain than their own since birth and have never interacted with rats of their own strain, they do not help rats of their own strain when tested as adults. Here we interrogated whether a cross-fostered rat expressed his lack of motivation to help through any behaviors beyond not-helping. Accordingly, a cross-fostered rat was placed in an arena with a trapped rat and the door to the centrally located restrainer was taped shut. We found that cross-fostered rats moved more slowly and approached the trapped rat less than did control, regularly-raised rats tested under the same conditions. We then asked whether the behavior of the cross-fostered rats influenced the trapped rat. After being restrained with cross-fostered rats, trapped rats showed a decrease in exploratory behavior in an open field test compared to trapped rats who were raised normally. The same decrease in movement was observed after subject rats were allowed to freely interact with cross-fostered rats. These results suggest that rats that do not help demonstrate their disinterest to a trapped rat and that trapped rats exposed to apathetic rats show behavior suggestive of an increase in anxiety. In sum, the paradigm introduced here could serve as a rodent model for social rejection.

## Introduction

Helping behaviors in rats have been reported in several experimental paradigms (Ben-Ami Bartal et al., 2011; Sato et al., 2015; Marquez et al., 2015; Hernandez-Lallement et al., 2015). In the restrainer paradigm, the potential helper rat (“free rat”) is placed in an arena with a rat trapped (“trapped rat”) in a restrainer that can only be opened from outside (Ben-Ami Bartal et al., 2011). We have previously demonstrated that an important factor in the free rat’s decision to help or not is whether the trapped rat is of a strain that the free rat has previous lived with (Ben-Ami Bartal et al., 2014). If the free rat has lived with even just one rat of the same strain as the trapped rat, then the free rat will help strangers of the cagemate’s strain. On the other hand, if the free rat has never lived with a rat of a given strain, the free rat will not help trapped rats of this unfamiliar strain. Remarkably, this effect holds even when the free rat is of the same strain as the trapped rat, as occurs when rats are fostered from birth with a foreign strain. Thus, Sprague-Dawley (SD) rats cross-fostered by Long-Evans (LE) rats from the day of birth, and never exposed to other SD rats prior to testing, do not help SD rats as adults but help LE rats in the restrainer paradigm (Ben-Ami Bartal et al., 2014).

In this study, we turned our attention from factors that influence the free rat’s motivation to help to the trapped rat’s response. In particular, we were interested in the effects of the lack of concern on the part of the cross-fostered SD (cfSD) rats. In other words, how does a trapped rat, who does not get helped by a member of his own strain, react to the free rat’s lack of concern, which we term “apathy” (Wieser et al., 2015).

We hypothesized that free rats behaviorally signal their social motivation even in the absence of an opportunity to help. In other words, we posit that a cfSD rat communicates his apathetic attitude toward a same-strain rat in distress through behaviors beyond the explicit fact of not-helping. We there placed a cfSD rat in an arena with a regularly-raised SD (rrSD) rat trapped within a restrainer that was taped shut. Analysis of this experiment showed that cfSD rats do indeed manifest disinterest in trapped SD rats. Therefore, we then asked whether the lack of prosocial signals from the free rat has affective consequences for the trapped rat. We predicted that a lack of demonstrative concern would lead to an affective reaction on the part of the trapped rat, the target of the free rat’s putative apathy. Accordingly, subject rats, who were always rrSD rats, were exposed to an “actor” rat, which was either a cfSD or a rrSD rat. In the first experiment, the subject rat was trapped for the duration of the exposure session while the actor rat was the free rat outside. In the second experiment the subject was free to interact with the actor rat in an open arena. Following each brief exposure, subject rats were then tested in an open field.

## Design and Results

### Experiment 1

A subject rat was placed in a restrainer that was located at the center of the arena (see Fig. 1B & 1C for set-up). Subject rats were rrSD rats and were neither kin nor familiar to any of the free rats used. The restrainer door was sealed closed to prevent helping, potential behaviors were limited to more nuanced demonstrations of interest and caring. Immediately after the interaction session, the subject rat was retrieved and tested in an open field test.

**Figure 1.**
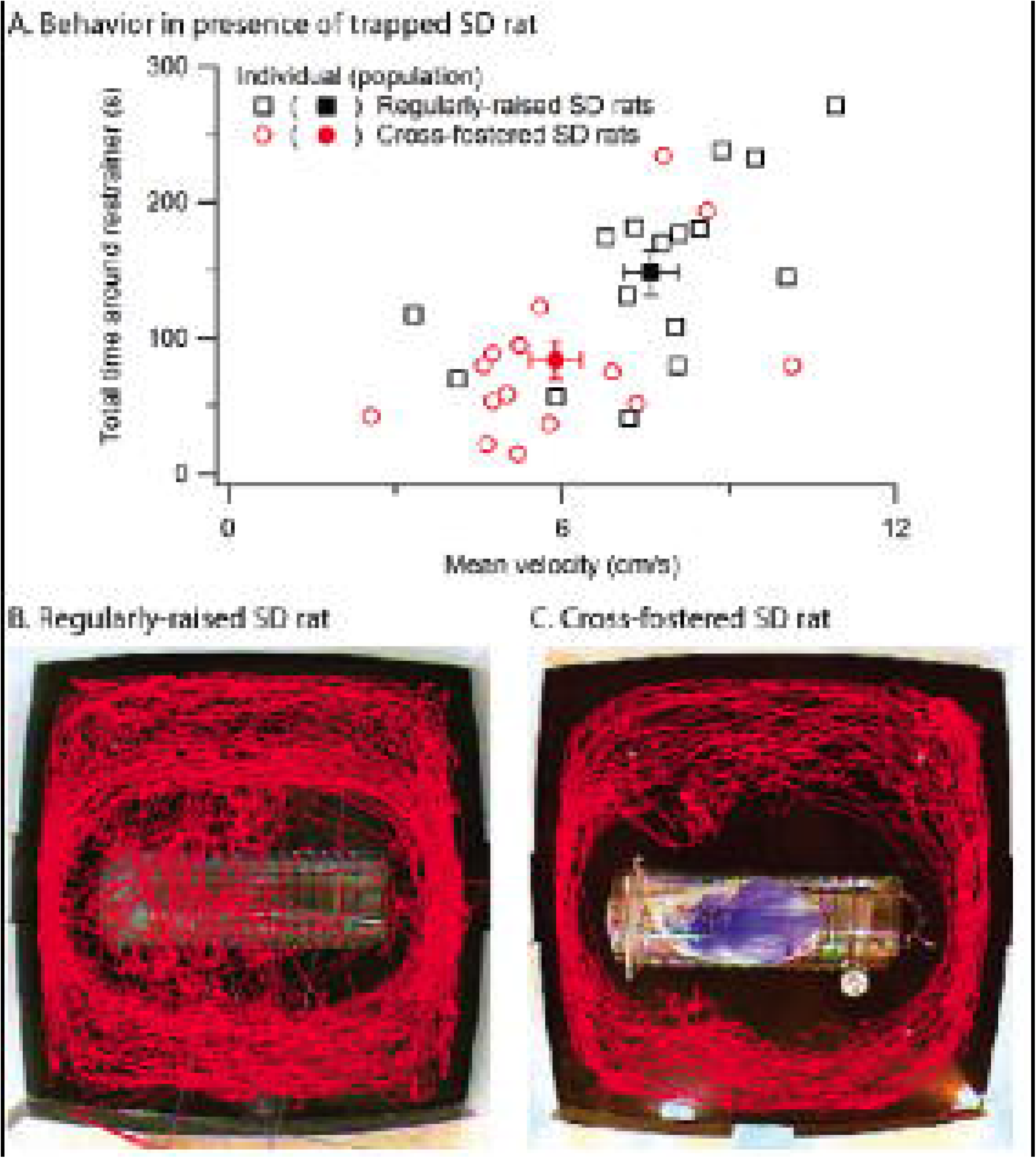
Apathy of cfSD rat is behaviorally manifest. A. Cross-fostered rats (circles) spent significantly less time around the restrainer, where the restrained conspecific was held; they also moved more slowly during the session than did rrSD rats (squares). Each open symbol represents one rat and the filled symbols represent the population averages (with SEM bars). B-C: Sample traces of the movements of a rrSD rat (B) and a cfSD rat (C) during the arena session are illustrated. Cross-fostered free rats spent less time around the restrainer and also moved less in general.

On the first day of testing, half of the subject group (N=8) were trapped with cfSD free rats and the other half (N=8) were trapped in the presence of free rrSD rats. Assignments were reversed on a second test day to ensure a counter-balanced design that enabled the statistical removal of individual differences between baseline activity levels of individual subject rats.

We first compared the movement of the free rat (cfSD vs. rrSD) during the interaction session (see methods). Two measurements, the free rat’s average velocity during the session and the cumulative time the free rat spent near the restrainer, were analyzed with a general linear model with two factors, *treatment* (cfSD vs. rrSD) and *day* (test day 1 vs. test day 2, accounting for potential order/repetition effects). The model revealed that cfSD rats had significantly slower movement (F(1.29)=8.0, Bonferroni p=0.02, Fig. 1A) and spent significantly less time around the restrainer (F(1, 29)=8.5, p-value=0.01, after Bonferroni correction, Fig. 1A). Figure 1B-C show sample traces of a rrSD rat and a cfSD rat during the arena session. These results suggest that cfSD rats had less interest in approaching and exploring the restrainer than did rrSD rats.

To compare activity levels of the subject rats in the two groups, we analyzed the open field results (average velocity, cumulative time spent in the center and cumulative time spent in corners; Table 1) with a general linear model. The model had three factors, *treatment* (cfSD vs. rrSD), *day* (test day 1 vs. test day 2, accounting for potential order/repetition effects) and *individual* (16 levels, one for each subject rat, accounting for predisposed differences between individual activity levels). Rats moved at a significantly faster speed on the second day of testing (F(1,14)=21.1, Bonferroni p<0.001). The model also revealed that subject rats moved at a lower average velocity after being trapped with cfSD free rats than with rrSD free rats (F(1,14)=11.3, Bonferroni p-value=0.015). However, there were no significant differences in cumulative time spent in the center or time spent in corners (Bonferroni p-value >1).

**Table 1.** Mean (± SEM) values for subject rats in Experiment 1.

| Subject rat measure during open field test | After interaction with regularly-raised actor rats | After interaction with cross-fostered actor rats |
| --- | --- | --- |
| Average velocity (cm/s) | 4.8 ± 0.3 | 4.2 ± 0.2 |
| Time in corners (s) | 185.4 ± 17.3 | 180.7 ± 15.4 |
| Time in center (s) | 38.8 ± 4.4 | 39.0 ± 5.5 |

#### Causal relationship between Free and Subject Rats’ Behaviors

In order to determine if a causal connection between the free rat’s lack of interest in the restrainer and the trapped rat’s subsequent demonstration of reduced mobility exists, we asked whether the restrainer-approach behavior of the free rat could explain the reduced mobility of the trapped rat *on an individual level*. We therefore pooled all open field velocity data (from cfSD and rrSD treatments) and analyzed them with a general linear model with three factors, *day* (test day 1 vs. test day 2), *individual* (16 levels, one for each subject rat) and *restrainer-approach time of the free rat* (covariate scale factor). Treatment was not included as a factor because differences in restrainer-approach time reflect the significant difference between treatments described above. The model revealed that the restrainer-approach time of the free rat has considerable explanatory power for the velocity of the subject rat in open field (F(1,14)=3.347, p=0.09).

#### Cage-front approach (CFA) analysis

We hypothesized that the difference in time spent around the restrainer by cfSD and rrSD rats is a socially specific effect and does not represent general differences in anxiety or locomotion. To test this, we compared the latency of approach by cfSD and rrSD rats to a non-social stimulus.

In the cage-front approach (CFA) test, the tops of housing cages were opened half-way (front half exposed) and the latency until a rat put his front paws on the front wall was recorded as the CFA score (see Methods). Thus, a higher CFA score indicates a longer period of time before the rat approached the novel stimulus, namely, the open part of the cage. A comparison revealed that cfSD rats had significantly lower CFA scores (p=0.002, unpaired two-tailed t-test) than did rrSD rats, indicating that cfSD rats moved more quickly toward the cage front than did rrSD rats. Yet in the presence of a trapped rat, cfSD rats moved more slowly on average than did rrSD rats These results suggest that the difference in the restrainer-centered behavior shown by rrSD and cfSD rats is specifically social.

### Experiment 2

To test whether interaction with cfSD rats creates behavioral differences even when the subject rat is not in distress (i.e. is not trapped in the restrainer), we modified Experiment 1 by removing the restrainer altogether, thereby allowing the two rats to interact freely. The arena session was reduced to 20 minutes. As was observed in Experiment 1, subject rats were significantly slower in open field after interacting with cfSD rats than with rrSD rats (F(1,14)=16.5, Bonferroni p =0.003; Table 2) and there was no significance difference in the two cumulative time measurements (corner F(1,14)=0.000, center F(1,14)=2.8 both Bonferroni p>1; Table 2).

**Table 2.** Mean (± SEM) values for subject rats in Experiment 2.

| Subject rat measure during open field test | After interaction with regularly-raised actor rats | After interaction with cross-fostered actor rats |
| --- | --- | --- |
| Average velocity (cm/s) | $5.2 \pm 0.4$ | $4.3 \pm 0.3$ |
| Time in corners (s) | $272.3 \pm 33.4$ | $272.7 \pm 28.8$ |
| Time in center (s) | $26.3 \pm 2.9$ | $23.4 \pm 3.5$ |

#### Anogenital sniffing analysis shows no observed difference in social activity

Frequency of anogenital sniffing is an established measure of social activity of rats (e.g., Witt et al., 1992). To test whether the behavioral difference in subject rats after having interacted with rrSD rats and cfSD rats is related to differences in the general sociability of rrSD and cfSD rats, we manually counted the number of anogenital sniffing events during interaction sessions (data not shown). A paired t-test revealed no significant differences related to treatment in the frequency of sniffing or being sniffed, or in the total number of sniffings and ratio between sniffing and being sniffed (all p-values above 0.4). Fighting took place occasionally in both groups, with the same incidence (4/16 sessions in either group).

## Discussion

Here we found that cfSD rats show their disinterest in distressed rrSD rats through lower activity and less approach. In support of our hypothesis, the differences in social behavior between cfSD and rrSD rats had a demonstrable effect on subject rats. Rats were less active in an open field test after exposure to a cfSD rat than after exposure to an rrSD rat. This difference was observed regardless of whether the subject rat was trapped and in distress or was allowed to roam about the arena.

The lower activity level and decrease in approach behavior shown by cfSD rats toward trapped rrSD rats may reflect a motoric difference, i.e. cfSD rats are generally slower than rrSD rats. This notion, however, is countered by the result of the CFA analysis, which shows that cfSD rats were *quicker* to approach an opening. Thus, the findings that cfSD rats were more active in their home cages and yet approached a trapped rat less is best explained by a lack of interest in the trapped rat. This idea is supported by the finding that cfSD rats do not show pro-social behavior toward rrSD rats even as they help rats from the foster strain (Ben-Ami Bartal et al., 2014). Moreover, trapped rrSD rats sense and react to the cfSD rat’s behavior in a manner consistent with being the target of an uncaring attitude.

After interacting with a cfSD rat, subject rats were less exploratory in an open field test. No measure of anogenital sniffing or fighting behavior differed between treatments, evidence that differences in the sociality or aggression of interactions are unlikely explanations for the differences in the subject rats’ exploratory behavior. On the other hand, an inverse correlation between restrainer-approach time of the free rat and velocity in open field of the subject rat suggests that reduced restrainer-approach time accounts for at least some of the behavioral effects on subject rats. Therefore, it is likely that subject rats are sensitive to cues from conspecifics that indicate their degree of empathic concern. Sensing the uncaring attitude of cfSD rats alters the behavior of targeted conspecifics, rendering them less exploratory. This is consistent with previous findings that rats exhibit decreased exploratory activity after enduring social stress (Scoifo et al., 2002).

The paradigm introduced here may serve as a rodent model of social rejection. It has advantages over the resident-intruder and pup separation tests, models of negative social interactions in adults and young, respectively (Hammels et al., 2015; Panksepp & Beatty, 1980). While socially defeated animals have a stress response similar to that exhibited by socially rejected humans, social defeat paradigms typically involve at least one direct instance of physical injury, whereas social rejection can be induced in humans with subtle, non-violent cues such as exclusion from a ball-toss game (e.g. Eisenberger et al., 2003). Separating pups from mothers carries immediate threats to survival, a contingency that does not exist in social rejection (Panksepp & Beatty, 1980). As is the case for human social rejection, our paradigm relies on subtle, non-violent cues, and does not threaten violence. That rats are affectively altered by a nuanced expression of disinterest suggests that social rejection may operate in rat societies as it does among humans. A rodent model for social rejection opens up possibilities for exploring the biological contributions to negative interactions in mammalian societies.

## Detailed Method

### Subjects

Male Sprague-Dawley rats (N=24, 2-month-old at the start of the study, Charles River, Portage, Michigan) were purchased for the regularly-raised Sprague-Dawley group (“rrSD”, N=8) and the subject SD group (“subject”, N=16). 8 timed-pregnant Sprague-Dawley rats and 8 timed-pregnant Long-Evans rats were purchased (Charles River, Portage, Michigan) to breed the male cross-fostered Sprague-Dawley rats (“cfSD”, N=8). The cfSD rats were pair-housed with an LE cagemate under the same conditions.

Adult male animals were pair-housed in a 12:12 light-dark cycle and given *ad libitum* access to food and water. Purchased male rats were given 2 weeks to acclimate to the housing environment. Pregnant rats were single-housed under the same conditions prior to birth.

### Cross-fostering Protocol

Pregnant rats were timed to given birth on the same day (*±1* day). Two male pups from each SD litter were taken on the day of birth and moved into a LE litter. After a week one pup from each litter was sacrificed. Pups were weaned two weeks after birth and each cfSD rat was pair-housed with an LE rat from the same litter.

### Setup

8 Plexiglas arenas (50 *×* 50 cm, 32–60 cm high) were used for the experiment. Restrainers (Plexiglas, 25 *×* 8.75 *×* 7.5 cm, Harvard Apparatus, Holliston, Massachusetts) were placed in the center of the arenas. Positioning of the restrainers were standardized across arenas and test days with painted marks on the arena floor. Restrainers had small openings that allowed tactile and olfactory communication. Restrainers were closed with customized doors (Plexiglas). In each session, the doors were taped shut to prevent any opening.

### Video Analysis

A CCD color camera (KT&C co., Seoul, South Korea) was mounted on each arena. Videos were analyzed with Ethovision (Noldus Information Technology, Inc. Leesburg, VA). Before analyzing, the frame rate of the videos was lowered to 15 fps to expedite processing. To enable tracking, free rats in the experiments were colored red with markers before every session, and subject rats were colored blue.

In the analysis of sessions with restrainers, “near the restrainer” was defined as a 25*35 cm rectangular area centered on the center of the restrainer. In the analysis of open field videos, “center” was defined as a 30 *×* 30 cm square in the middle of the arena and “corner” was defined as the difference between the arena and the arena’s inscribed circle.

